# IL-17A potentiates malignant T-cell viability via electron transport chain complex I in cutaneous T-cell lymphoma

**DOI:** 10.1101/2025.05.21.655309

**Authors:** Diksha Attrish, Bhavuk Dhamija, Ditipriya Mukherjee, Vinanti Sawant, Soumitra Marathe, Dominik Saul, Moumita Basu, Ankit Banik, Neha Sharma, Tanuja Shet, Hasmukh Jain, Robyn Laura Kosinsky, Priyanka Sharma, Rahul Purwar

## Abstract

Cutaneous T-cell lymphoma (CTCL) is a rare malignancy of skin-resident T cells characterized by an aberrant cytokine profile, notably elevated Interleukin-17A (IL-17A) levels. However, the functional importance of IL-17A dysregulation in the CTCL pathogenesis remains poorly understood. Our analysis revealed increased IL-17A receptor (IL-17RA) expression in CTCL patient samples, supporting the importance of IL-17A signaling in CTCL progression. Proteomic and metabolomic profiling of IL-17A-treated malignant T cells revealed mitochondrial electron transport chain (ETC) complex I as a key downstream target of IL-17A signaling. Furthermore, we confirmed the elevated expression of ETC complex I in CTCL patient samples compared to healthy control T cells, hence highlighting the clinical relevance of the IL-17A signaling-ETC complex 1 axis. Altogether, our data unveil the function of IL-17A signaling in the metabolic axis in CTCL progression, which could be useful for developing advanced therapeutic strategies.

**<u>Significance statement:</u>** We uncover an IL-17A-induced electron transport chain (ETC) complex I pro-tumorigenic circuit and provide proof of concept evidence that ETC complex I can be targeted to combat the challenges posed by Cutaneous T-cell Lymphoma aggressiveness.

## Introduction

Cutaneous T-cell lymphoma (CTCL) is a heterogeneous group of malignancies arising from skin-homing malignant T-cells with diverse clinical, histological, and molecular features (1,2). The two primary subtypes of CTCL are Mycosis fungoides (MF), the most common subtype, and Sézary syndrome (SS). MF is characterized by the proliferation of skin-resident effector memory T-cells with cerebriform nuclei. Whereas, SS is characterized by circulating central memory T-cells, erythroderma, and lymphadenopathy, representing advanced-stage disease (3,4). In its early stages, MF remains confined to the skin; however, around 20% of MF cases progress to advanced stages that involve lymph nodes and organs, which are associated with high mortality (5,6). Patients presenting with extracutaneous spread of MF/SS experience markedly poorer outcomes than those without and have five-year survival rates as low as 24% (7). A key feature of advanced-stage MF/SS is the transition of small-to-medium-sized malignant T-cells into large, blast-like T-cells, a process known as large cell transformation, which is associated with poor survival (8). Immune dysregulation plays a vital role in the pathogenesis of lymphoma. As the disease progresses, the tumor microenvironment undergoes significant changes, including increased levels of Th2 (T helper 2) cytokines, Th17 cells (producing IL-17A among other members of the family), and regulatory T cells (9).

IL-17A is a proinflammatory cytokine essential for immune defense, and its dysregulation contributes to chronic inflammation in several diseases (10–12). IL-17A transmits the signal by binding to a heterodimeric receptor complex formed by IL-17 receptor A (IL-17RA) and IL-17 receptor C (IL-17RC) (13). In CTCL patients, IL-17RA levels are notably elevated, whereas IL-17RC levels remain unchanged (14). Furthermore, elevated levels of IL-17A and its receptor, IL-17RA, are both conspicuously increased and are known to be associated with CTCL progression (14,15). Therefore, elevated IL-17RA levels raise the possibility of IL-17A signaling and its functional relevance in CTCL, which has not been investigated yet.

Besides its primary roles in immune regulation, emerging evidence suggests that IL-17A also plays critical roles in metabolic processes (16). In intestinal epithelial cells, IL-17A modulates metabolic pathways to regulate cell proliferation, facilitating tissue repair (17). In melanocytes, restraint stress induced IL-17A, leading to mitochondrial dysfunction (18). In severe asthmatic bronchial fibroblasts, IL-17A promotes autophagy-mediated mitochondrial dysfunction (19). We recently showed that the IL-17A signaling induces the metabolic transcription factor Hypoxia-inducible factor 1-alpha (HIF-1α), enhancing glucose, glutamine, and fatty acid metabolism, ultimately driving keratinocyte hyperproliferation (20). Moreover, IL-17 signaling is known to promote metabolic reprogramming in activated fibroblastic reticular cells, which enhances glucose uptake and upregulates key enzymes involved in oxidative phosphorylation (OXPHOS). Mitochondrial electron transport chain (ETC) complexes, particularly complexes I and IV, play a pivotal role in cancer metabolism by supporting OXPHOS, maintaining mitochondrial membrane potential (MMP), and regulating reactive oxygen species (ROS). ETC complex I is notably upregulated in acute myeloid leukemia cells (21), while the overexpression of MTCO1 (mitochondrially encoded cytochrome C oxidase 1), a component of complex IV, is linked to enhanced proliferation in breast and bladder cancers (22,23). These findings suggest a broader role for ETC activity in tumorigenesis, with emerging evidence implicating IL-17 signaling in the regulation of mitochondrial bioenergetics.

Even though metabolic reprogramming is a hallmark of cancer cells, allowing cancer cells to survive under energy-restricted conditions, the role of IL-17A in neoplastic T-cell metabolism remains largely unexplored. Activated normal T lymphocytes and neoplastic T lymphocytes exhibit overlapping yet distinct metabolic rewiring, presenting a potential therapeutic window for cancer treatment. Tumor cells often enhance the OXPHOS biogenesis to sustain their growth despite metabolic constraints (24–26). Understanding how IL-17A influences neoplastic T-cell metabolism could uncover novel therapeutic strategies for targeting metabolic vulnerabilities in cancer.

Using high-throughput proteomics and nuclear magnetic resonance (NMR) spectroscopy, we identified ETC complex I along with MTCOI as a downstream target of IL-17A signaling. Subsequent validation experiments confirmed that both ETC complex I and MTCO1 are consistently regulated by IL-17A across multiple human CTCL cell lines. We observed elevated ETC complex I levels in CTCL patients in comparison to healthy samples, suggesting its role as a critical metabolic adaptation for tumor growth. Biochemical assays showed that IL-17A enhances ETC complex I activity, increasing NAD⁺/NADH and ATP/ADP ratios, and hyperpolarizing mitochondria. Pharmacological inhibition of ETC complex I using mubritinib reversed IL-17A-driven metabolic rewiring and significantly reduced cell proliferation. Together, our findings position ETC complex I as a therapeutic vulnerability in IL-17A-driven CTCL.

## MATERIALS AND METHODS

### Patient CTCL Biospecimens

Punch skin biopsies, as well as blood samples from CTCL patients (age range: 19-59, n = 14, Table 1) with known blood involvement, were collected from Tata Memorial Hospital following protocols approved by the Institutional Ethics Committee of Tata Memorial Centre and Indian Institute of Technology (IIT) Bombay, under documents numbers IITB-IEC/2016/017. It was also ensured that patients were treatment naïve (except for topical steroids). Tumors were classified according to the Tumor, Node, Metastasis, Blood staging criteria. Patients with co-existing skin ailments such as psoriasis or autoimmune disorders or viral infections (including HIV/HCV/HBV), or those on immunosuppressive therapy, were excluded from the study. Healthy individuals were recruited based on the absence of any disease presentation, inflammatory skin disorders, and infections. Informed consent was obtained from all subjects, and all studies were conducted in accordance with both the Declaration of Helsinki and Ethics Committee guidelines.

**Table 1.** Patient sample information. Details of enrolled patients with CTCL and healthy control samples used in this study. CTCL: cutaneous T-cell lymphoma; MF: mycosis fungoides.

| Sample | Age | Sex | Type of Disease | Stage |
| --- | --- | --- | --- | --- |
| T-cells isolated from CTCL patients (Figure 1 and 3) |  |  |  |  |
| 1 | 48 | M | MF | Stage IIA |
| 2 | 58 | M | MF | Stage IV |
| 3 | 23 | F | CTCL | Stage IIB |
| 4 | 47 | M | MF | Stage III |
| 5 | 57 | M | MF | Stage IV |
| 6 | 81 | M | MF | Unavailable |
| 7 | 45 | M | CTCL | Stage IIA |
| 8 | 41 | F | MF | Stage III |
| 9 | 43 | M | CTCL | Stage IB |
| T-cells isolated from healthy donors (Figure 1 and 3) |  |  |  |  |
| 1 | 45 | M | Healthy | Not applicable |
| 2 | 42 | M | Healthy | Not applicable |
| 3 | 39 | M | Healthy | Not applicable |
| 4 | 31 | M | Healthy | Not applicable |
| 5 | 35 | F | Healthy | Not applicable |
| 6 | 32 | F | Healthy | Not applicable |
| 7 | 30 | M | Healthy | Not applicable |
| 9 | 28 | M | Healthy | Not applicable |
|  |  |  | Healthy | Not applicable |
| CTCL Skin biopsy (Figure 1 and 3) |  |  |  |  |
| 1 | 24 | M | MF | Stage IB |
| 2 | 60 | M | MF | Stage IIB |
| 3 | 59 | F | Primary cutaneous T-cell lymphoma | Stage IIB |
| 4 | 45 | M | MF | Stage IB |
| 5 | 66 | M | MF | Stage IA |
| Healthy control skin biopsy (Figure 1 and 3) |  |  |  |  |
| 1 | 53 | F | Healthy | Not applicable |
| 2 | 40 | M | Healthy | Not applicable |
| 3 | 49 | F | Healthy | Not applicable |
| 4 | 37 | F | Healthy | Not applicable |
| 5 | 27 | M | Healthy | Not applicable |

### Cell Culture

Human T-cell lymphoma cell lines, Jurkat E6.1(RRID: CVCL_0065), Hut-78 (RRID: CVCL_0337), and MJ (RRID: CVCL_1414) were obtained from ATCC (Manassas, VA). Cells were cultured in RPMI 1640 (Jurkat) medium supplemented with 10% fetal bovine serum (FBS) or with 20% FBS (HuT 78) or Iscove’s Modified Dulbecco Medium (MJ) 20% FBS supplemented with 1% penicillin-streptomycin (Gibco, Waltham, MA, USA). All the cell lines were tested negative for mycoplasma contamination (G238, Applied Biological Materials Inc.). To isolate PBMCs, whole blood was mixed with an equal volume of PBS and layered over Ficoll and centrifuged at 23°C for 30 minutes. The resultant buffy coat, a distinct layer, was collected and washed with PBS two times, and peripheral blood mononuclear cells (PBMCs) were collected.

T-cells were separated from PBMCs by immunogenic negative selection using the EasySepTM Direct Human T-cell Isolation Kit (STEMCELL Technologies, Cambridge, MA, USA, 19661). Briefly, PBMCs were resuspended in PBS. Cells were incubated with an antibody cocktail (50 μl/ml) and beads (50 μl/ml) for 5 minutes at room temperature (RT). Cells were placed in the magnet, and T-cells were collected by pipetting out the supernatant. To ensure the maximum purity, cells were again incubated with beads (50 μl/ml) and placed in a magnet for 5 minutes at RT. The supernatant was collected, and this process was repeated to ensure optimal bead removal. Once isolated, the cells were quantified and activated with CD3/CD28 Dynabeads Human T-cell activator (1:1 ratio of beads to cells; Gibco, 11131D) with IL-2 (50 ng/ml) in AIM V media with 10% heat-inactivated FBS and 1% penicillin. After 48 hours, beads were removed, and cells were subsequently cultured with IL-2 (20 ng/ml).

### Antibodies and Reagents

The following antibodies were used for immunofluorescence (all from ThermoFisher Scientific): anti-human CD3 (CD3-12; MA5-16622, RRID: AB_2538118), anti-human MTCO1 (clone 1D6E1A8, 4596000, RRID: AB_10374492), anti-human IL17RA (PA5-34571, RRID: AB_2551923), anti-human complex I (clone 18G12BC2, 43-8800, RRID: AB_2533543), anti-NDUFS2 (GeneTex, GTX04733) and following antibodies were used for flow cytometric analyses (all from ThermoFisher Scientific): Alexa Fluor APC-conjugated anti-CD217 (IL17Ra, 424LTS, 17-7917-42, RRID: AB_10597595) IgGκ1 isotype control (17-4714-42, RRID: AB_1603315), Alexa Fluor 488-conjugated goat anti-mouse IgG (H+L) cross-adsorbed (A11001, RRID: AB_2534069), Alexa Fluor 647-conjugated donkey anti-rat IgG (H+L) highly cross-adsorbed (A78947, RRID: AB_2910635) and Alexa Fluor 594 conjugated anti-rabbit (A11037, RRID: AB_2534095), Alexa Fluor 488 conjugated anti-rabbit (A11008, RRID: AB_143165), Alexa Fluor conjugated anti-rat 594 (A11007, RRID: AB_10561522) Following kits were used (Sigma-Aldrich): ADP/ATP ratio assay kit (MAK135), Mitochondrial Isolation kit (MITOISO2), Mitochondrial Complex I activity assay kit (MAK359), Easy direct human T cell isolation kit (STEM cell, 19661), Dynabeads human T-cell activator (STEMCELL, 11131D). Following staining, dyes were used: MitoSOX™ Mitochondrial Superoxide Indicators (Invitrogen, M36008), JC-1 Dye (Mitochondrial Membrane Potential Probe, Invitrogen, T3168), MitoTracker™ Red CMXRos (Invitrogen A66443). Mubritinib (Sigma-Aldrich SML1312), HClO_4_ Merck (109065), K_2_CO_3_ (Merck 104928).

### Immunohistochemistry Staining

The tissue sections were collected in 10% Neutral buffered formalin. The tissues were dehydrated using alcohol gradients, followed by incubation in xylene and melted paraffin wax. Tissue blocks were prepared, and 5 µm sections were immobilized on silane-coated slides (BioMarq, India). Slides were dried overnight at room temperature. For staining of formalin-fixed paraffin-embedded tissues, the slides were preheated at 60℃ for an hour and incubated in xylene two times for 5 minutes each. The tissues were rehydrated by a series of alcohol gradients: 100%, 95%, 90%, and 85%, followed by one incubation in distilled water, each incubation lasting for 5 minutes. Antigens were retrieved by heating the slides in citrate buffer (10 mM sodium citrate, 0.05% Tween 20, pH 6.0; Himedia and Merck). Sections were blocked with 5% BSA (HiMedia) for 60 minutes and incubated overnight at 4°C with anti-CD3 antibodies (1:100), anti-MTCO1 (1:400), anti-IL-17RA (1:200), and anti-NDUFS2 (1:17) diluted in blocking solution containing 0.1% Triton X-100 (Merck). The sections were incubated with Alexa Fluor 488 or Alexa Fluor 594-conjugated secondary antibodies (1:500) and counterstained with 4′,6-diamidino-2-phenylindole (DAPI), dried, mounted with ProLong Gold Antifade Mountant (ThermoFisher Scientific). Sections were imaged with an LSM 780 microscope (Zeiss, 20 Oberkochen, Germany) under 20x and 63x magnification using ZEN Black software (Zeiss, 21 Oberkochen, Germany). The mean fluorescent intensity (MFI) was measured with ImageJ software (27).

For immunofluorescence staining of cells, 0.1 million cells were seeded per well in an 8-well chamber plate (IBIDI) with or without IL-17A treatment. After 24 hours, cells were washed with PBS and incubated with 4% paraformaldehyde (HiMedia) for 20 minutes. Cells were permeabilized with 0.1% Triton 100 (Merck) for 30 minutes and blocked with 2% BSA for 60 minutes. Cells were incubated with anti-complex I (1:1000) and anti-MTCO1 (1:500) antibody for 2 hours at RT, followed by 60 minutes of incubation with Alexa Fluor 488 goat anti-mouse antibody. Further, cells were counterstained with DAPI for 5 minutes and imaged using an EVOS M7000 Imaging System (ThermoFisher Scientific). ImageJ software was used to measure the mean intensity of the images.

### Single-cell data analysis

Transcriptome-wide analysis of CD3+ lymphocytes from skin biopsies of patients (n=5: CTCL2, CTCL5, CTCL6, CTCL8, CTCL12) with advanced-stage CTCL and 4 healthy donors (n=4: HC1, HC2, HC3, HC4) at a single cell level was based on a previously published dataset (28). Here, a 10X Chromium Platform was used along with the STAR RNA-seq aligner on the human genome GRCh38, and processed along the Cell Ranger pipeline (10x Genomics). Data with at least 200 unique molecular identifiers (UMIs) per cell were extracted, normalized, and integrated using the Seurat package v5.0 in R4.2.0. After quality control and integration, we performed a modularity-optimized Louvain clustering. A subsequent R package was ggplot2 (3.5.0.9000).

### Sample Preparation for high-throughput label-free quantification using LC-MS/MS

Jurkat cells were seeded in 12-well plates and stimulated with IL-17A (50 ng/ml) for 24 hours at 37℃ with 5% CO2. Samples were processed using the Trizol-chloroform method, and proteins were isolated. The pellet was washed with 3M Guanidine Hydrochloride solution in 95% ethanol and sonicated, followed by 100% ethanol washes. The protein pellet was resolubilized in the Urea-Thiourea buffer (UTB). The protein extract (35 μg) was reduced using dithiothreitol (10 mM) for 60 minutes at 50℃, followed by alkylation with iodoacetamide (15 mM) for 30 minutes at RT in the dark. The volume of the digestion mixture was increased by 4x using 50 mM ammonium bicarbonate, and protein was digested using 1μg MS-grade trypsin. The peptides thus obtained were cleaned using C18-ziptips and quantified using Scopes’ method. Subsequently, 2 μg of clean peptide samples was injected into the Thermo Scientific Q-Exactive Plus through the EasyNLC HPLC autosampler and C18 PepMap TM trap column.

### Proteomics data primary analysis and label-free quantification

The MS spectra were analyzed using the Thermo-scientific mass informatics platform, Proteome Discoverer version 2.2. Workflows for discovery proteomics were used with both Mascot and SequestHT as search engines. For label-free quantification, the standard LFQ workflows by Thermo were used. The samples were normalized concerning the peak area based on the “total peptide amount”. The permitted retention time shift was set to 2 minutes. Fragment mass tolerance was set to 0.02 Da, and the precursor mass tolerance to 2 ppm. The false discovery rate (FDR) was set to 0.01 (Strict). Proteins were filtered based on applying stringent, high-confidence filters (unique peptide ≥ 2), protein fold change ≥ 1.2 upregulation, and abundance ratio P ≤ 0.1. Importantly, unique peptides were considered for peak-area-based quantification. Null hypothesis testing was performed using ANOVA (background-based) to analyze identified proteins statistically.

### 1H NMR Sample Preparation and Metabolic Flux Computational Analyses

Jurkat cells were stimulated with IL-17A (50 ng/ml). Cells were collected after 24 hours, and metabolites were isolated using a methanol-based extraction protocol (29). Samples were vacuum-dried and stored at -80°C. 1H NMR was performed on a Bruker Spectrometer operating at 750.0 MHz at a temperature of 298 K. NMR spectra were analyzed using Chenomx NMR Suite 8.3, and concentrations of metabolites were calculated using DSS as an internal standard. Metabolic flux modeling was performed using manually curated, constraint-based human metabolic network software (MitoCore) as described (30,31). The starting point was to calculate the flux of the central metabolites to be used as input for MitoCore model (**see supplementary method**). This led us to simulate the metabolism by using flux balance analysis (FBA), where the model calculated the flux of metabolites through a network of biochemical reactions, assuming maximum ATP production as the objective function.

### Flow cytometry

To assess IL-17RA expression, cells were incubated with APC-conjugated anti-IL-17RA and respective isotype control for 30 minutes. Samples were analyzed on a BD FACSVerse system (BD Biosciences, NJ, USA) using BD FACSSuite software.

For intracellular staining, cells (0.1 million/well) were seeded into a 24-well plate and treated with IL-17A. After 24 hours, cells were extracted and washed using PBS. Cells were fixed with 80% methanol (100 μl per sample) for 5 minutes at RT, washed with wash buffer, and treated with 0.1% Tween-20 for 20 minutes at RT. Cells were again washed with wash buffer and incubated in 0.3 M glycine (100 μl per sample) for 30 minutes at RT. Cells were washed and permeabilized with the permeabilization buffer (BD FACS Perm Wash Buffer). The cells were incubated with the anti-complex I and anti-MTCO1 primary antibodies for 30 minutes. Cells were washed with permeabilization buffer and stained with Alexa Fluor 488 goat anti-mouse antibody (1:300). Finally, cells were washed, and data were acquired by BD FACSVerse flow cytometer (BD Biosciences).

### Enzymatic Activity Assays

Mitochondria were purified using a mitochondrial isolation kit (Merck, MITOISO2). ETC complex I activity was measured using a complex I activity kit according to the manufacturer’s instructions (Merck).

### NAD^+^/NADH Assay

NAD^+^ concentration: Cells were washed and incubated in 0.5 M HClO_4_ (Merck). Cells were neutralized with 0.55 M K_2_CO_3_ (Merck), and cell extract was collected. NAD^+^ (Merck) standards were prepared, ranging from 50 µM to 500 µM. Standards and Samples were loaded and incubated in NAD^+^ assay buffer (lactate & 1U/ml LDH in Tris-glycine-hydrazine, pH 9) for 60 minutes at 37°C. Absorbance was recorded at 340 nm.

NADH concentration: Cells were washed and incubated in NADH extraction buffer (50 mM NaOH and 1 mM EDTA) for 30 minutes at 60℃. The reaction was neutralized with NADH Neutralization buffer (0.3 M Potassium phosphate buffer, pH 7.4), and the supernatant was collected. NADH (Merck) standards ranged from 100 nM to 100 µM, and samples were loaded in a 96-well plate (CoStar, Corning, NY, USA). Absorbance was recorded at 340 nm.

### ADP/ATP Assay

Cells were seeded and treated with IL-17A and mubritinib (200 nM, Merck). The luminometric assay was performed according to the instructions given in the ADP/ATP assay kit (Merck).

### Mitochondrial Oxidative Stress (reactive oxygen species) Evaluation

Cells were treated with IL-17A and mubritinib. Samples were stained with 5 µM MitoSOX Red (ThermoFisher Scientific) in 200 μl PBS for 20 minutes. Data was acquired on BD FACSVerse (BD Biosciences) instrument.

### Mitochondrial Membrane Potential Evaluation

Cells (0.1 million/well) were seeded and treated with or without IL-17A. Cells were collected, and 50 μM hydrogen peroxide was added in the positive control, followed by incubation at 37℃ for 5 minutes. Cells were washed with PBS and stained with 100 nM of Mitotracker and incubated for 30 minutes at 37°C. For staining with JC-1, cells were stained with 1 µM of JC-1 for 20 minutes at 37°C. Data was acquired on a BD FACSVerse (BD Biosciences) instrument.

### Apoptosis Assay

Cells were treated with the cytokine IL-17A for 24 hours. Cells were washed with Annexin V binding buffer (1X) and stained with PE-conjugated Annexin V and 7-AAD (Annexin V Apoptosis Detection Kit, Thermo Fisher Scientific). Flow cytometric analyses for early- and late-stage apoptosis and necrosis were performed on a BD FACSVerse (BD Biosciences) instrument.

### Cell Viability Assay

Cells (0.01 million/well) were seeded and cultured for 24, 48, and 72 hours with or without IL-17A or mubritnib (50-1000 nM) or DMSO. Post-incubation cells were harvested and stained with 0.4% trypan blue dye and cell viability was calculated.

### Statistical Analysis

An unpaired, two-tailed Student t-test was used to compare data in the Figures. 1 and 3-5. Background-based ANOVA was used to calculate the significance of protein or peptide changes in Figure 2. Prism 8.0.2 software (GraphPad, La Jolla, CA, USA) was used for statistical analysis.

**Figure 1.**
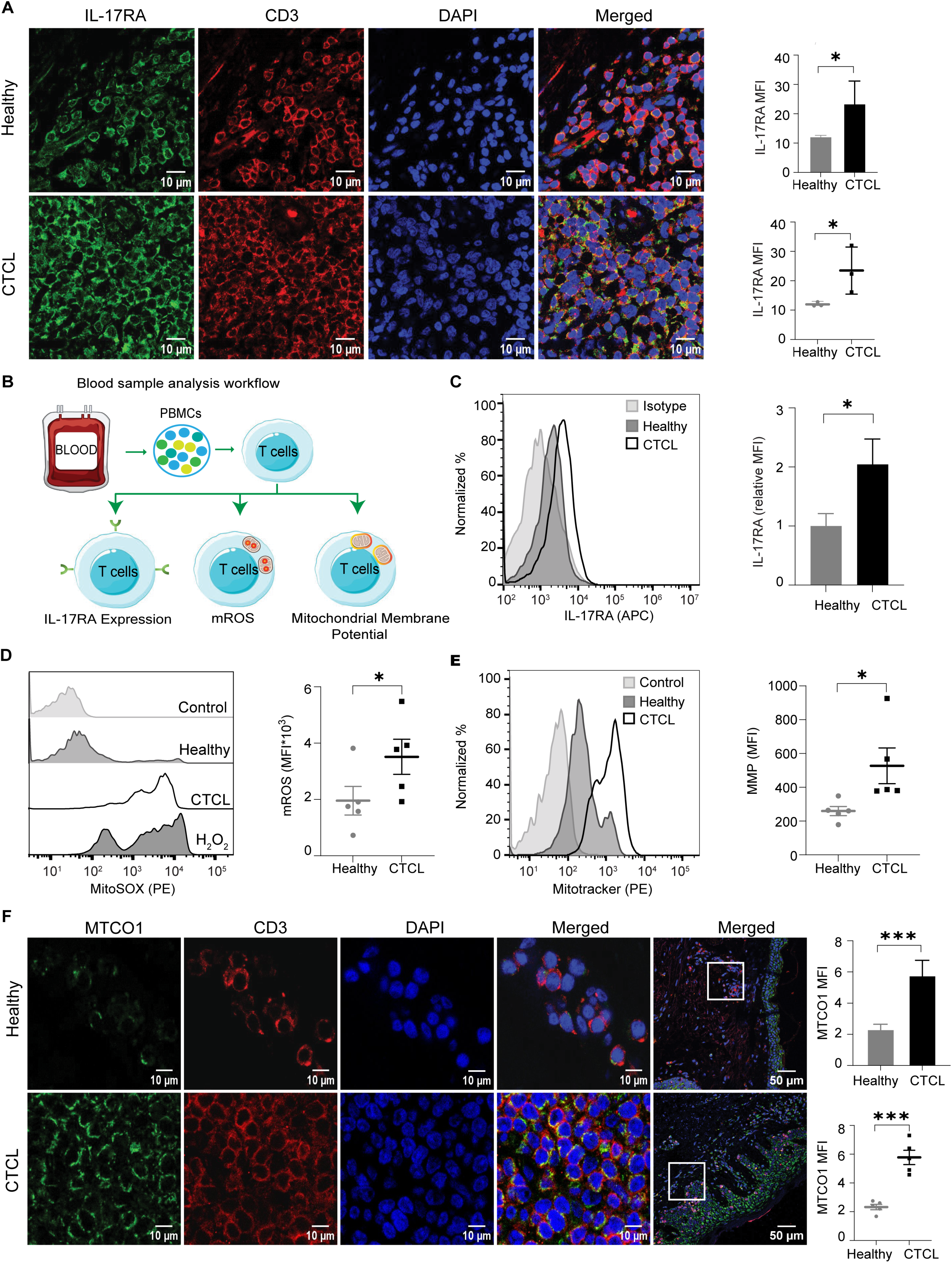
(Related to Figure S1). Elevated IL-17RA expression and mitochondrial metabolic anomalies in CTCL. (**A**) Representative immunofluorescence images (left) and quantification (right) of healthy versus CTCL patient skin biopsies for IL-17RA (green), CD3 (red), and DAPI (blue) with merged image. Bar plot showing mean fluorescence intensity of IL17RA for healthy and CTCL skin biopsies. Scatter plot showing data of individual samples (n=3). (**B**) Workflow of patient blood sample analysis. (**C**) Representative histogram and quantification of IL-17RA levels (n = 5) in T-cells isolated from CTCL blood samples and healthy donors. (**D-E**) Representative histogram and quantification of mitochondrial ROS (mROS, n=5) (**D**) and MMP (n = 5) **(E)** in T-cells acquired from CTCL patients enriched for malignant T-cells and healthy donors. **(F)** Representative images (left) and quantifications (right) of immunofluorescence staining of skin biopsy samples from healthy and CTCL patients showing MTCO1 (green), CD3 (red), and DAPI (blue) with merged images at two resolutions. Bar plot represents the mean fluorescence intensity of MTCO1 in healthy versus CTCL skin biopsies. Dot plot illustrates data from individual samples (n=5). All data are represented as mean ± SD. Staining intensity was measured using Image J software and is expressed as mean fluorescent intensity (MFI). Values are expressed as mean ± SEM. *p < 0.05, **p < 0.01, ***p < 0.001 (Student unpaired test). CTCL: Cutaneous T-cell Lymphoma, H2O2: Hydrogen peroxide, APC: Allophycocyanin, PE: Phycoerythrin.

**Figure 2.**
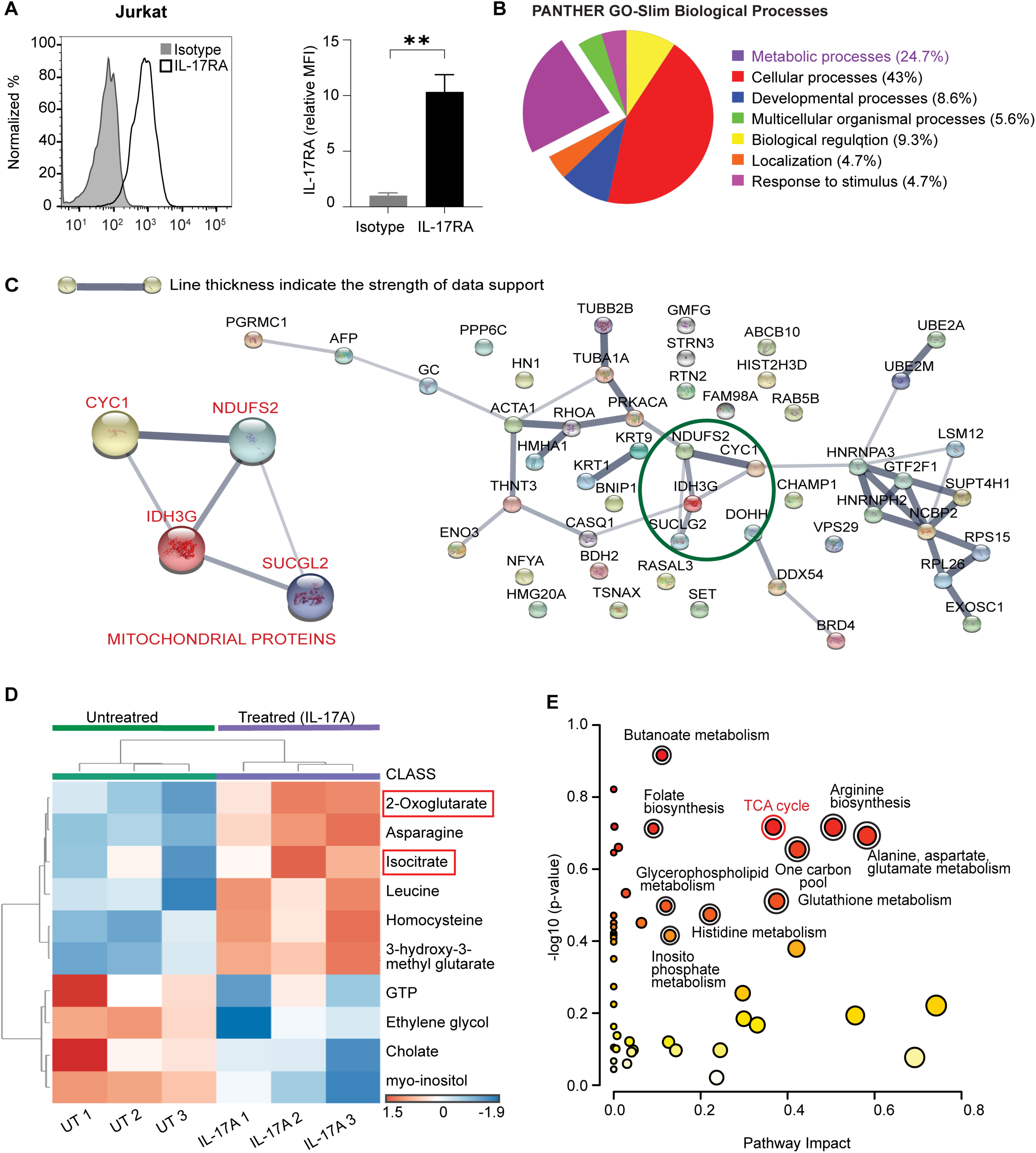
(Related to Figure S2, Table 2 and Table S1-S2). Characterization of metabolic pathways dysregulated due to IL-17A signaling by LC/MS-MS and NMR analysis. (**A**) Representative histogram and statistical representation quantifying IL-17RA expression on Jurkat cells. (**B**) Pie chart depicting functional pathways altered in IL–17A–treated versus untreated cells (n = 3) based on frequencies of differentially expressed (p < 0.1) proteins as determined using PantherDB software. **(C)** STRING network for all differentially expressed proteins in response to IL-17A treatment. **(D**) Heatmap of significantly upregulated metabolites in response to IL-17A treatment versus untreated cells (n=3) as determined by metaboanalyst. **(E)** Pathway analysis. The horizontal coordinate indicates the pathway impact value, the vertical coordinate indicates the −log10 (p-value) of the pathway, a dot in the Figure represents a metabolic pathway, the size of the dot is proportional to its impact value, and the colour of the dot represents the size of the pathway P value, where the colour change from yellow to red represents the change in the p-value from largest to smallest.

**Figure 3.**
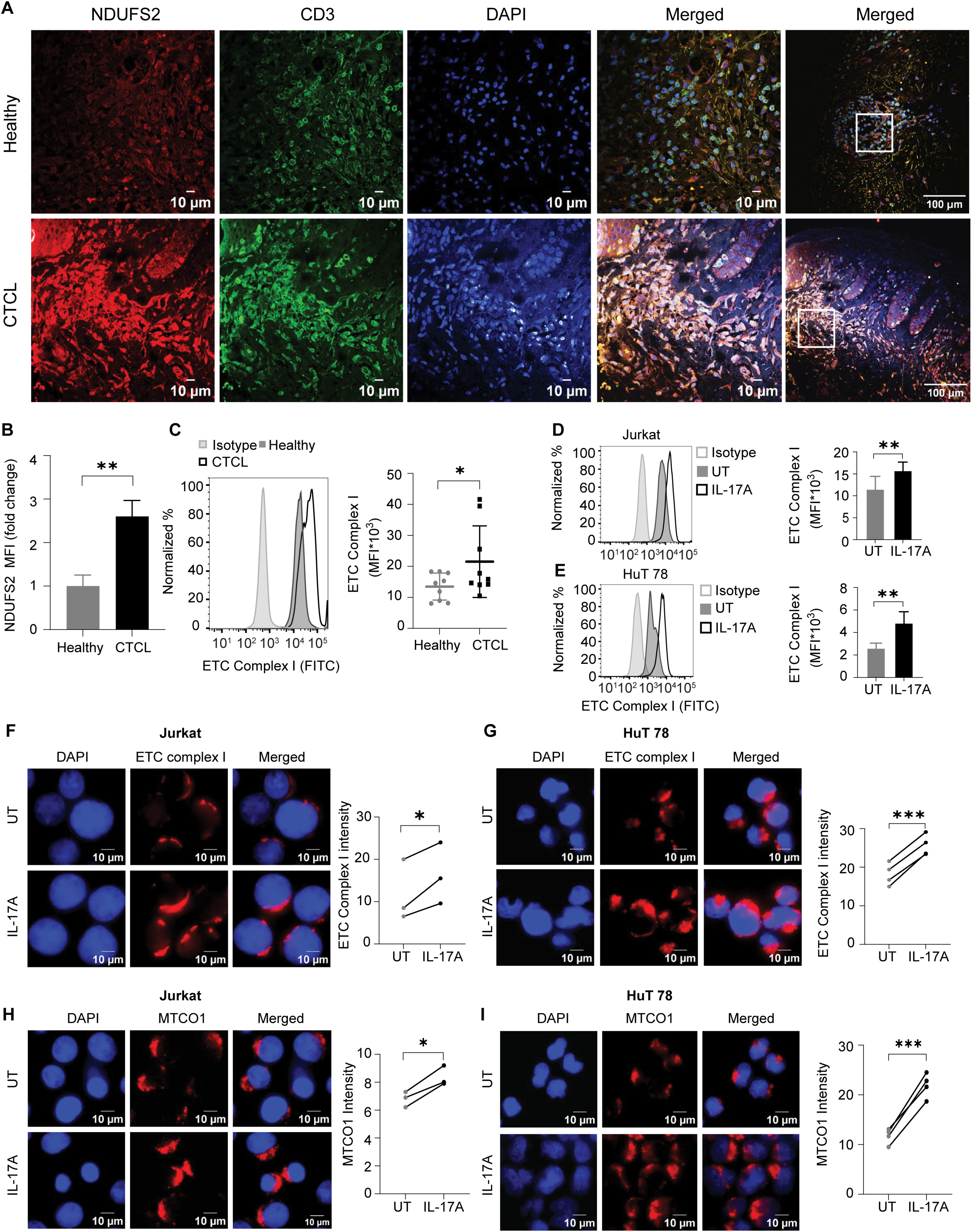
(Related to Figure S3 and Table 2) OXPHOS proteins are highly expressed in CTCL clinical samples and IL-17A-treated lymphocytes. **(A)** Representative immunofluorescence staining of healthy versus CTCL patient skin biopsies for NDUFS2 (red), CD3 (green), and DAPI (blue) with merged images at different indicated resolutions **(B)** Bar plot showing mean fluorescence intensity of NDUFS2 for healthy and CTCL skin biopsies (n=5). **(C)** Representative histogram and scatter plot (mean ± SD) quantifying ETC complex I expression in T-cells isolated from CTCL blood sample and healthy donors (n=9). (**D**-**E**) Cytometry histograms (left) and quantifications (right, MFI (MFI ± SD) of ETC complex I protein levels in (**D**) Jurkat and (**E**) HuT 78 cells. (**F**-**G**). Representative immunocytochemistry staining of ETC complex I (red), nucleus (DAPI) in (**F**) Jurkat and (**G**) Hut78 cells. (**H**-**I**). Immunocytochemistry staining showing MTCO1(red), nucleus (DAPI) in (**H**) Jurkat and (**I**) HuT 78 cells. Values are expressed as mean ± SEM. *p < 0.05, **p < 0.01, ***p < 0.001. UT: untreated, FITC: Fluorescein isothiocyanate.

**Table 2.**
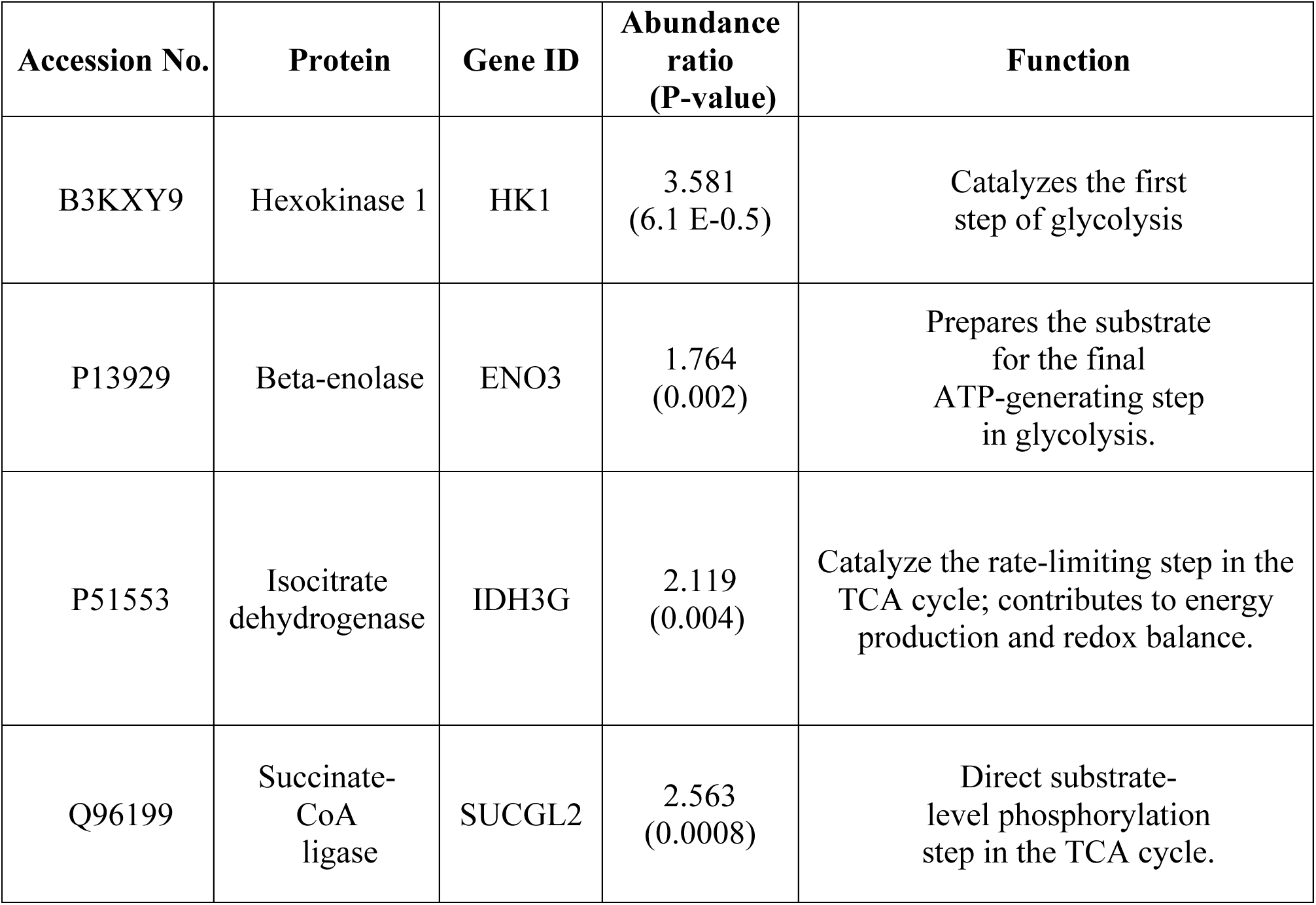
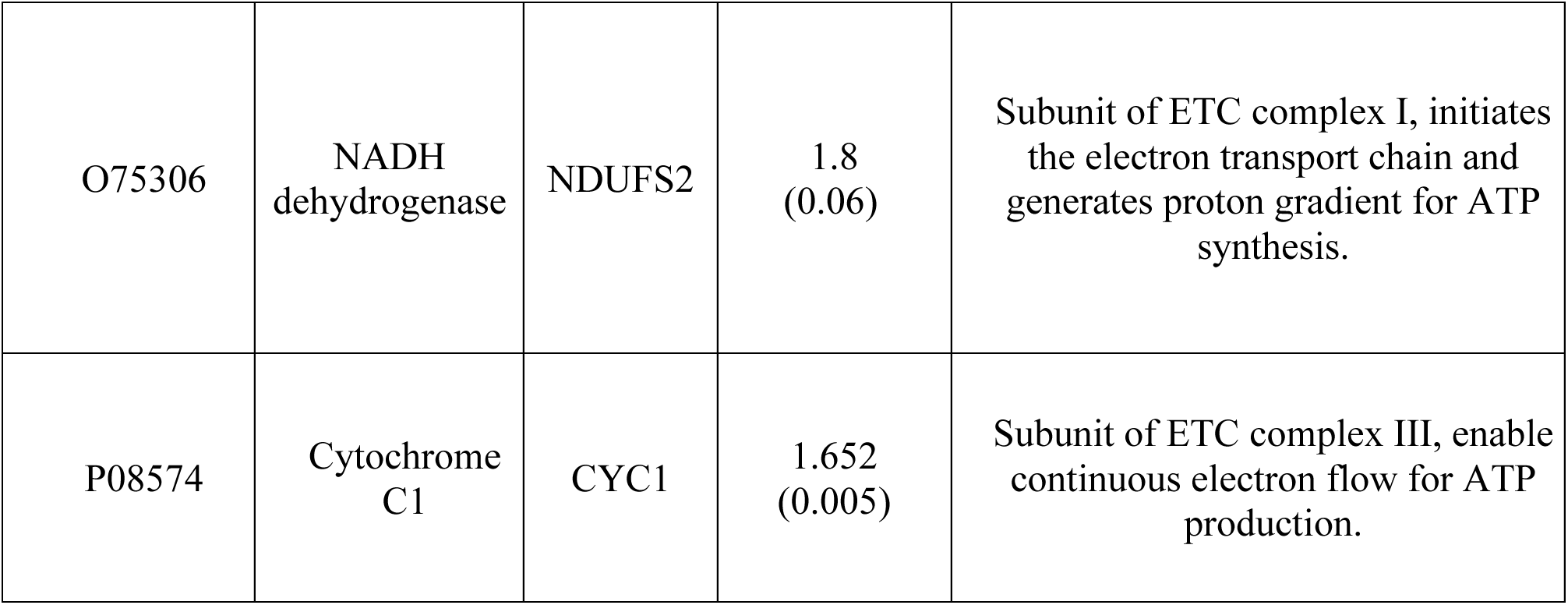
Differential expression of metabolic proteins. Annotated list of differentially expressed metabolic proteins in Jurkat T-cell lysate treated with or without IL-17A as determined by LC-MS/MS.

| Accession No. | Protein | Gene ID | Abundance ratio (P-value) | Function |
| --- | --- | --- | --- | --- |
| B3KXY9 | Hexokinase 1 | HK1 | 3.581<br>(6.1 E-0.5) | Catalyzes the first step of glycolysis |
| P13929 | Beta-enolase | ENO3 | 1.764<br>(0.002) | Prepares the substrate for the final ATP-generating step in glycolysis. |
| P51553 | Isocitrate dehydrogenase | IDH3G | 2.119<br>(0.004) | Catalyze the rate-limiting step in the TCA cycle; contributes to energy production and redox balance. |
| Q96199 | Succinate-CoA ligase | SUCGL2 | 2.563<br>(0.0008) | Direct substrate-level phosphorylation step in the TCA cycle. |
| O75306 | NADH dehydrogenase | NDUFS2 | 1.8<br>(0.06) | Subunit of ETC complex I, initiates the electron transport chain and generates proton gradient for ATP synthesis. |
| P08574 | Cytochrome C1 | CYC1 | 1.652<br>(0.005) | Subunit of ETC complex III, enable continuous electron flow for ATP production. |

### Data Availability

Data generated in this study can be requested from the corresponding authors Rahul Purwar and Priyanka Sharma. The raw data from high-throughput proteomics experiments of Jurkat cells are stored in the MassIVE knowledgebase, an online public repository for mass spectrometric data, under the specified doi:10.25345/C5M902F58

## RESULTS

### Blood samples and skin biopsies of CTCL patients show increased IL-17A receptor expression and enhanced mitochondrial metabolism

To determine if IL-17A induced signaling through IL-17RA in CTCL, we quantified the IL-17RA level in skin biopsies of CTCL patients and healthy controls using immunofluorescence. We found significantly higher IL-17RA levels in CD3+ lymphocytes infiltrating CTCL skin lesions than in those of healthy controls (23 ± 8 vs. 12.26 ± 0.72, p = 0.037, n=3) **(Figure 1A)**, confirming elevated IL-17A receptor expression in CTCL patients. Next, we examined IL-17RA expression in T-cells in the blood of CTCL patients (**Figure 1B**). Indeed, T-cells isolated from the blood of CTCL patients exhibited 2-fold higher IL-17RA levels than in T cells from the blood of healthy controls (p = 0.0208, n=5) **(Figure 1C)**, hence validating an induced IL-17A signaling in the CTCL patients. Additionally, we analyzed an RNA sequencing dataset (GSE128531) (28) and found that IL-17RA was highly expressed in CTCL patient samples compared to healthy controls **(Figure S1A-B)**. Collectively, these results illustrate the increased expression of IL-17RA on T-cells from both the malignant lesional skin and blood of CTCL, supporting its potential functional role in disease pathology.

Consistent with the established role of IL-17A in regulating the mitochondrial metabolism, we observed higher levels of mROS (3878 ± 625.65 vs. 2313 ± 547.4, p = 0.0442, n=5, **Figure 1D**) and MMP (749.2 ± 264.4 vs. 400 ± 266.7, p = 0.0198, n=5, **Figure 1E**) in CTCL samples than in healthy controls. To further investigate mitochondrial bioenergetics, we analyzed the expression level of MTCO1 and found a significantly higher MTCO1 level in CD3+ lymphocytes from CTCL skin specimens than in those of healthy skin (5.151 ± 0.53 vs. 2.326 ± 0.186, p = 0.0008, n = 5, **Figure 1F**). Our findings suggested a functional role for IL-17A in driving metabolic adaptations in the CTCL progression.

### IL-17A dysregulates the tricarboxylic acid (TCA) cycle and the OXPHOS pathway

To examine whether the observed enriched expression of IL-17RA on malignant CTCL T-cells functionally regulates metabolism, we performed a comprehensive analysis in Jurkat cells (an immortalized human T lymphocyte cell line) as they expressed high IL-17RA (**Figure 2A**). Besides this, activated T-cells, which are marked by high expression of CD25 **(Figure S2A),** also tested positive for IL-17RA expression **(Figure S2B)**. Jurkat cells exhibited IL-17RA expression levels comparable to activated T cells, and hence, we used them as our primary model system. To assess whether this is a generalized phenomenon, we also measured the IL-17RA expression in HuT 78 (a SS cell line, **Figure S2C**) and MJ (an MF cell line, **Figure S2D**). To assess the impact of IL-17A on the cellular metabolism, we performed high-throughput label-free quantitative liquid chromatography-tandem mass spectrometry (LC-MS/MS) using proteins isolated from Jurkat cells without (control) or with IL-17A treatment identified 337 significantly differentially expressed proteins (**Table S1**). Functional annotation and pathway analysis using the Panther DB tool revealed that 24.7% of proteins upregulated in IL-17A-treated cells belonged to cellular metabolic pathways (**Figure 2B**). Importantly, among the metabolic proteins, we observed an increase level of hexokinase 1 (HK1) and beta-enolase (ENO3) involved in glycolysis, isocitrate dehydrogenase (IDH3G) and succinate CoA-ligase (SUCGL2) in the TCA cycle, as well as NADH dehydrogenase (NDUFS2), a mitochondrial ETC complex I subunit, and cytochrome C1 (CYC1), a mitochondrial ETC complex III subunit, essential for OXPHOS (**Table 2)**. Additional analysis by the STRING pathway identified significant enrichment of mitochondrial proteins, including NDUFS2, CYC1, SUCGL2, and IDH3G (**Figure 2C**). These results thus demonstrated the notable upregulation of TCA cycle and OXPHOS proteins in response to IL-17A treatment in Jurkat cells.

Next, we performed the 1D-NMR metabolic profiling to monitor the metabolites in Jurkat cells altered by the IL-17A treatment (**Table S2**). TCA metabolites, including 2-oxoglutarate and isocitrate, were highly upregulated (**Figure 2D**), and pathways involved in the TCA cycle, glutamine/glutamate, and aspartate metabolism were altered (**Figure 2E**) in response to IL-17A treatment. Using all the differential metabolites obtained from the NMR (**Table S2**), the MitoCore simulation tool was used to calculate the metabolic flux (**see supplementary method**) of IL-17A-dysregulated reactions. Metabolic flux refers to the rate at which metabolites are utilized in biochemical reactions and is primarily regulated by enzyme activity. Analysis of metabolic fluxes (**Table 3**) revealed enhanced consumption of glutamine and increased synthesis of glutamate, as reflected by the elevated flux through glutaminase (GLS). The resulting glutamate was subsequently converted to 2-oxoglutarate, as evidenced by high flux through glutamate dehydrogenase (GDH). In addition to GDH activity, increased flux through isocitrate dehydrogenase (IDH) further contributed to 2-oxoglutarate synthesis. The accumulation of 2-oxoglutarate supported anaplerotic input into the TCA cycle, as indicated by enhanced flux through succinyl-CoA ligase (SUCGL) (**Table 3**). Moreover, the metabolic model predicted a concurrent increase in the flux through the ETC Complex I, linking enhanced mitochondrial oxidative metabolism with upregulated TCA cycle activity. Collectively, proteomics and metabolomics analysis demonstrated that IL-17A potentiated mitochondrial metabolic reprogramming in lymphocytes by enhancing the metabolic flux through the TCA cycle and the OXPHOS biogenesis **(Figure S2F)**. These observations confirmed the potential link between IL-17A and metabolic remodeling.

**Table 3.** Fold change (IL-17A-treated/untreated) in metabolic flux using the Mitocore simulation by performing flux balance analysis (FBA).

| Enzyme | Fold change in metabolic flux (p value) | Function |
| --- | --- | --- |
| Glutaminase (GLS) | 2.20<br>(0.0448) | Catalyze the conversion of glutamine to glutamate |
| Glutamate dehydrogenase (GDH) | 1.65<br>(0.0004) | Catalyze the conversion of glutamate to 2-oxoglutarate |
| Isocitrate dehydrogenase (IDH) | 5.05<br>(0.0311) | Catalyze the oxidative decarboxylation of isocitrate to synthesize 2-oxoglutarate in TCA cycle |
| Succinate Co-A ligase (SUCGL) | 1.62<br>(0.0433) | Convert succinyl-CoA to Succinate in TCA cycle |
| NADH dehydrogenase (ETC complex I) | 1.56<br>(0.0158) | Catalyze electron transfer from NADH to ubiquinone in OXPHOS pathway |

### IL-17A-treated lymphocytes and CTCL clinical samples highly express OXPHOS proteins

As analyses thus far indicated that IL-17A increased the metabolic flux through the TCA cycle and the OXPHOS machinery, we focused on the significantly upregulated core subunit of the ETC complex I NDUFS2 **(Figure 2C**, **Table 2)**. Quantitative analysis revealed a significant 2.6-fold increase (p = 0.0012, n=5) in NDUFS2 protein levels in skin biopsies from CTCL patients relative to healthy controls (**Figure 3A-B**). Analysis of published single-cell RNA sequencing data also demonstrated higher NDUFS2 expression in skin biopsies from CTCL patients than in those from healthy controls **(Figure S3A-B)**. Consistently, T cells derived from CTCL patients had a significantly higher level of ETC complex I than in those from healthy controls (21525 ± 3849 vs. 13496 ± 1456, n = 9) **(Figure 3C)**. These results confirmed the elevated levels of ETC Complex I, along with increased IL-17RA expression, in CTCL biospecimens compared to healthy controls.

To further strengthen that IL-17A could regulate the expression of ETC complex I, we performed the flow cytometry analysis in malignant T-cells with or without the IL-17A treatment. We found a significant increase in ETC complex I protein levels both in Jurkat cells (15757 ± 2105 vs. 11517 ± 3242, p =0.0084, n = 6, **Figure 3D)** and HuT 78 cells (4827 ± 1089 vs. 2610 ± 462.7, p = 0.0034, n = 6, **Figure 3E)**, hence confirming elevated ETC complex I level upon IL-17A induction. Similarly, we found that IL-17A treatment increased the ETC complex I expression in other IL-17RA-positive cells, including MJ (**Figure S3C**), and activated T-cells **(Figure S3D)**. Moreover, immunofluorescence labeling revealed a notable increase in ETC complex I protein in IL-17A-treated Jurkat **(Figure 3F)** and HuT 78 (**Figure 3G)** cells. This confirmed that IL-17A signaling positively regulated the ETC complex I expression. Next, we investigated IL-17A-mediated regulation of MTCO1, which was enriched in CTCL skin biopsies (**Figure 1F**). Immunofluorescence analysis demonstrated higher expression of MTCO1 in IL-17A-treated Jurkat **(Figure 3H)** and HuT 78 **(Figure 3I)** cells. In line with this, flow cytometry analysis revealed a substantial increase in MTCO1 in Jurkat **(Figure S3E)**, HuT 78 **(Figure S3F)**, and MJ **(Figure S3G)** cells. Moreover, flow cytometry analysis of activated T cells **(Figure S3H)** revealed enhanced MTCO1 levels when exposed to IL-17A. Thus, a prominent increase in proteins involved in OXPHOS indicated the biogenesis of OXPHOS machinery in T-cell lymphoma lines treated with IL-17A. These results showed that IL-17A signaling significantly impacted mitochondrial proteins in lymphoma cells.

### IL-17A enhances mitochondrial bioenergetics and promotes malignant cell growth

Next, we examined whether IL-17A signaling affects the metabolic processes related to ETC complex I activity. We first checked the effect of IL-17A on the ETC complex I enzymatic activity and found a substantial increase (p = 0.0127, n=4, **Figure 4A)** in Jurkat cells. Next, we studied the impact of IL-17A on the ATP production (32) and found a notable reduction (p = 0.045, n = 5) in the ADP/ATP ratio in Jurkat cells treated with IL-17 in comparison to the control **(Figure 4B)**, indicating that ATP pool was being generated and ADP being consumed at a higher rate. HuT 78 cells similarly displayed a consistent increase (p = 0.0006, n = 4) in ETC complex I activity **(Figure 4C)** and marked reduction (p = 0.006, n = 5) in the ADP/ATP ratio **(Figure 4D)** in IL-17 treated condition in comparison to control, thus strengthening the effect of IL-17A on the ETC complex I activity. We also examined the influence of IL-17A on NAD^+^ and NADH levels and found a significant increase in NAD^+^ levels (24.16 ± 1.28 vs. 19.57 ± 2.136, p = 0.0081, n = 6, **Figure 4E**) with a concomitant decrease in NADH concentration (24.48 ± 3.59, vs. 30.62 ± 7.3, p = 0.03, n = 4, **Figure 4F**) in Jurkat cells treated with IL-17A. Similarly, HuT 78 cells treated with IL-17A also displayed an increase in NAD^+^ levels (101.1 ± 3.8 vs. 75.81 ± 3, p = 0.0139, n = 3, **Figure 4G**) and a significant decrease in NADH levels (95.53 ± 18.74 vs. 121.2 ± 43, p = 0.0186, n = 4, **Figure 4H**). Thus, the increased ATP production and NAD^+^ levels strongly correlated with the enhanced ETC complex I activity.

**Figure 4.**
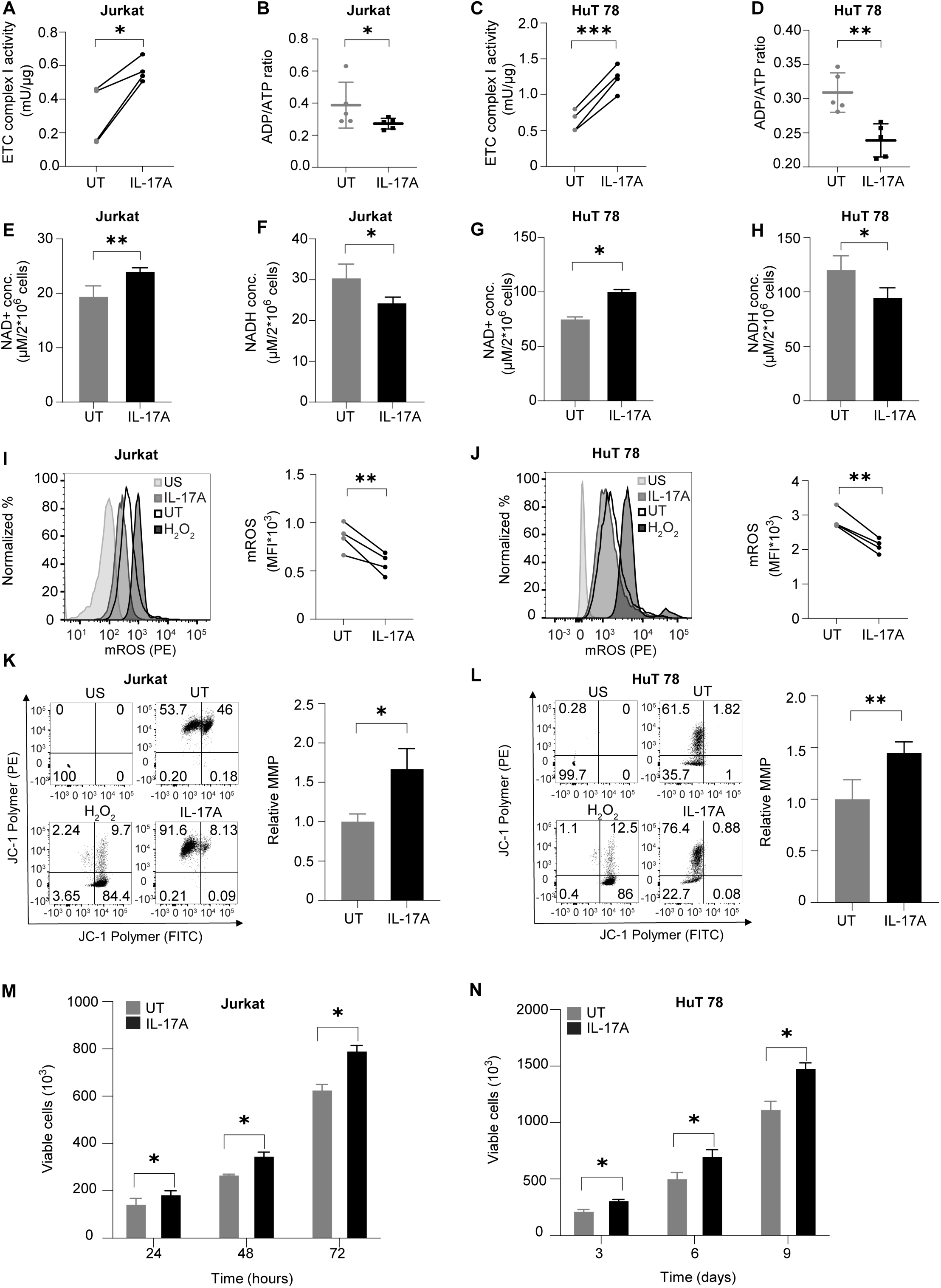
(Related to Figure S4) IL-17A rewires the mitochondrial metabolism and affects cellular physiology. **(A)** Enzymatic activity of ETC complex I in Jurkat cells. **(B)** Luminometric quantification of ADP/ATP ratio (n=5) in Jurkat cells. **(C)** Enzymatic activity of ETC complex I in HuT 78 cells. **(D)** Luminometric quantification of ADP/ATP ratio (n=5) in HuT 78 cells. **(E)** NAD^+^ levels and **(F)** NADH levels in Jurkat cells (n=3). **(G)** NAD^+^ levels and **(H)** NADH levels in HuT 78 cells (n=3). Representative histogram (left) and paired line plot (right) quantifying mROS (n=3) by flow cytometry in **(I)** Jurkat and **(J)** HuT 78 cells. Flow cytometry dot plots and statistical analysis representing mitochondrial membrane potential (MMP) using JC-1 (n=3) in **(K)** Jurkat and **(L)** HuT 78 cells. Cell viability measurement by trypan blue counting (n=3) in **(M)** Jurkat and **(N)** HuT 78 cells. Results represent n = 3-6 independent experiments, and values are expressed as mean ± SEM. *p < 0.05, **p < 0.01, ***p < 0.00. US: unstained, UT: untreated, PE: phycoerythrin.

In parallel, we characterized the effect of IL-17A on mitochondrial metabolism. MitoSox staining demonstrated a marked decrease in mROS levels in Jurkat **(Figure 4I)** and HuT 78 cells **(Figure 4J)** and JC-1 staining showed augmented MMP levels in both Jurkat (p = 0.0144, n = 4) and HuT 78 cells (p = 0.0061, n =3) cells **(Figure 4K-L)** upon IL-17A stimulation. Mitotracker staining also demonstrated higher MMP in Jurkat **(Figure S4A)** and HuT 78 cells **(Figure S4B)** treated with IL-17A, further confirming the hyperpolarized state of mitochondria in the IL-17A milieu. Subsequently, apoptosis assay revealed a significantly reduced apoptotic cell population in IL-17A-treated samples compared to untreated controls **(Figure S4C)**. Additionally, cell viability assays demonstrated that IL-17A treatment promoted robust proliferation in both Jurkat (**Figure 4M**) and HuT 78 (**Figure 4N**) cells. These findings collectively indicate that IL-17A-driven dysregulation of mitochondrial metabolism enhances cell survival in malignant lymphocytes.

### Inhibiting ETC complex I activity impairs mitochondrial metabolism and limits cell proliferation

Finally, we evaluated the functional impact of the pharmacological inhibition of ETC complex I in T-cell lymphoma cell lines using mubritinib (21). Treating cells with mubritinib reduced the ETC complex I activity significantly in Jurkat **(Figure 5A**, **Figure 5SA)** and HuT 78 also demonstrated a similar reduction in complex I activity (**Figure 5B**, **Figure 5SB**). We further studied the effect of ETC complex I inhibition on the ADP/ATP ratio, NAD^+^ and NADH levels, mROS and MMP. Mubritinib treatment increased the ADP/ATP ratio significantly in IL-17A-treated Jurkat cells **(Figure 5C)** as well as HuT 78 cells **(Figure 5D)**, respectively. Consistently, mubritinib-treated Jurkat cells and HuT 78 cells exhibited a substantial decrease in NAD^+^ levels **(Figure 5E-F)**, accompanied by a marked accumulation of NADH **(Figure 5G-H)**. MitoSOX staining revealed a significant accumulation of mROS in mubritinib-treated lymphoma cells (**Figure 5I–J**). Furthermore, mubritinib treatment led to significant reductions in MMP in IL-17A-treated Jurkat cells (p = 0.0033, n = 4; **Figure 5K**) and HuT 78 cells (p = 0.013, n = 3; **Figure 5SC**), respectively. Finally, we evaluated the effect of ETC complex I inhibition on the lymphoma cell viability. In Jurkat cells, mubritinib treatment significantly reduced the cell viability by 48% at 48 hours (p = 0.0029, n = 4) and 62% at 72 hours (p = 0.0007, n = 4) (**Figure 5L**). Similarly, the viability of HuT 78 cells treated with mubritinib, compared to untreated, decreased by 50% at 48 hours (p = 0.0014, n = 4) and 63% at 72 hours (p = 0.0001, n = 4) (**Figure 5M**). These data supported the role of ETC complex I as a key downstream mediator of IL-17A-induced cell proliferation. Moreover, pharmacological targeting of ETC complex I could be a promising metabolic vulnerability with the potential to reduce lymphoma cell viability and enhance therapeutic efficacy in CTCL.

**Figure 5.**
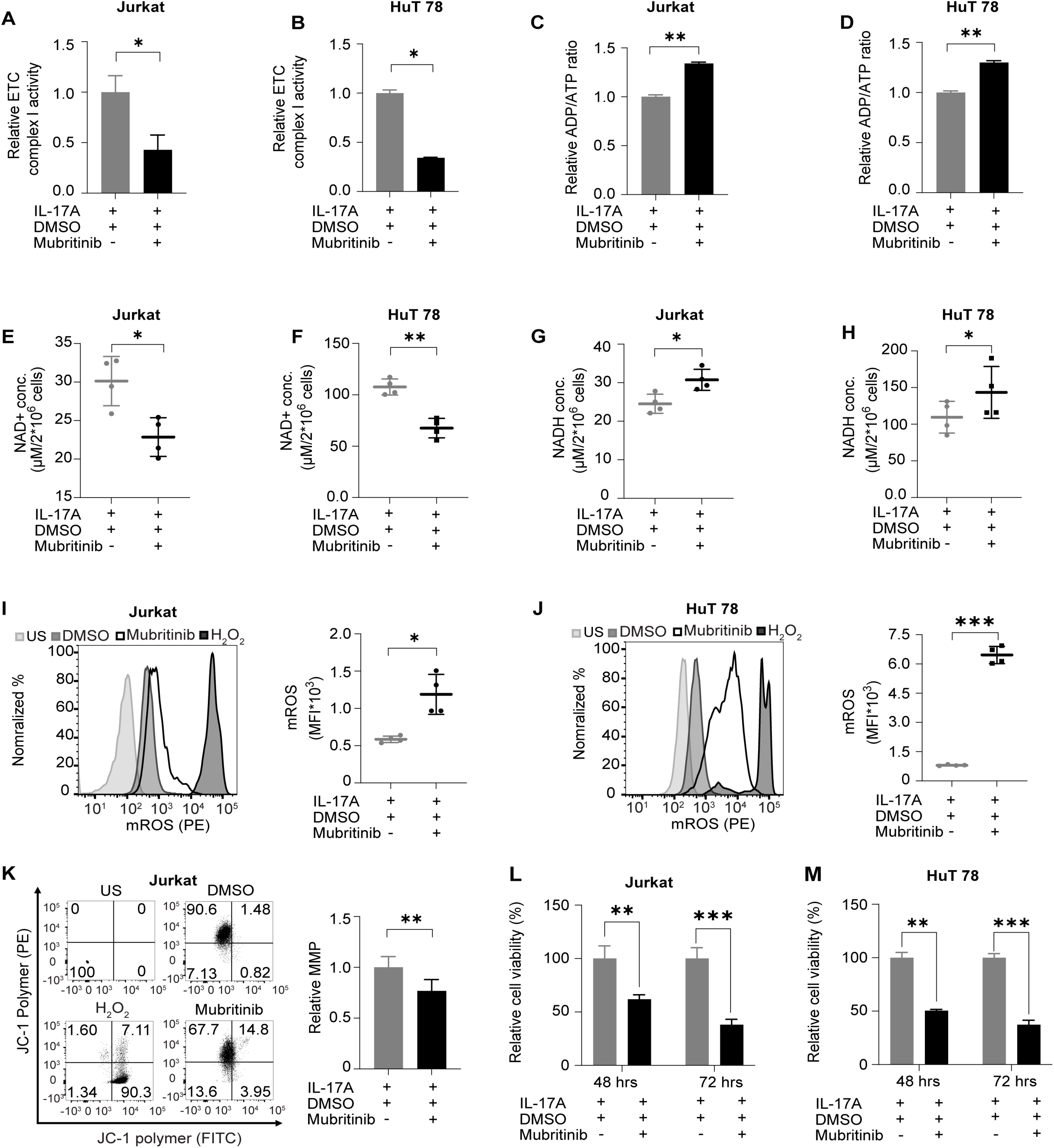
(Related to Figure S5) Inhibiting complex I perturbed IL-17A-mediated metabolic rewiring and induced oxidative stress and apoptosis. **(A)** Effect of mubritinib on the enzymatic activity of the ETC complex I in Jurkat cells (n = 5) and **(B)** HuT 78 cells (n=5). **(C-D)** Luminometric quantification of the ADP/ATP ratio upon mubritinib treatment in (**C**) Jurkat cells (n=5) and **(D)** HuT 78 cells (n=5). Effect of mubritinib on NAD^+^ levels in **(E)** Jurkat cells and **(F)** HuT 78 cells. Assessment of NADH levels in **(G)** Jurkat cells and **(H)** HuT 78 cells following mubritinib treatment. Flow cytometry histogram and scatter plot quantifying mROS upon mubritinib treatment (n=4) in **(I)** Jurkat and **(J)** HuT 78 cells. Flow cytometry dot plots and bar plot quantifying MMP upon mubritinib treatment (n=4) in **(K)** Jurkat cells. Effect of mubritinib on cell viability under IL-17A treatment in **(L)** Jurkat and **(M)** HuT 78 cells. Values are expressed as mean ± SEM. *p < 0.05, **p < 0.01, ***p < 0.001. US: unstained, UT: untreated, DMSO: dimethyl sulfoxide (vehicle control), PE: phycoerthrin. Mubritnib dose used for Jurkat and HuT 78 cells is 200 nM.

## DISCUSSION

The predominance of IL-17A within the CTCL tumor microenvironment raises an important question about its functional role in disease progression. While previous studies have implicated IL-17A in exacerbating the CTCL progression (15), our findings shed light on a previously underexplored dimension of IL-17A function and its involvement in metabolic reprogramming. Here, we systematically examined the IL-17A-mediated metabolic pathways in cancerous T-cells using multi-disciplinary approaches. Results of our immunofluorescence analysis using skin tissue biopsies of CTCL patients and healthy controls indicate active IL-17A signaling in the tumor niche. Transcriptomic profiling of advanced CTCL skin biopsies confirms elevated IL-17RA expression, which is independently supported by single-cell RNA sequencing data from a separate CTCL patient cohort (28). Importantly, our data reveal a metabolic dimension to this signaling pathway. Elevated mitochondrial ROS and enhanced MTCO1 expression in cancerous cell lines and patient samples suggest that IL-17A may promote mitochondrial bioenergetics. This aligns with the emerging understanding that immune and cancer cells often rely on mitochondrial function to support their proliferation and survival (25). In this context, IL-17A signaling may serve as a metabolic driver that enhances oxidative phosphorylation and supports the energetic and biosynthetic demands of malignant T cells. The concurrent upregulation of IL-17RA in both tissue and transcriptomic datasets, along with the observed metabolic alterations, underscores a potential link between inflammatory signaling and mitochondrial reprogramming in CTCL. These findings suggest that IL-17A may facilitate the disease progression not only through immune modulation but also by sustaining mitochondrial function and redox homeostasis in tumor cells.

Our multi-omics analysis reveals that IL-17A regulates key metabolic pathways, including carbohydrate metabolism, the TCA cycle, and oxidative phosphorylation, and implicates ETC complex I as a critical downstream target of IL-17A signaling. ETC complex I is essential for the regeneration of NAD⁺, which in turn sustains the continuity of the TCA cycle (33). Therefore, we hypothesized that elevated ETC complex I activity could enhance TCA cycle anaplerosis by maintaining NAD⁺ availability, while simultaneously supporting mitochondrial metabolism through increased ATP production and reactive oxygen species (ROS) generation. To validate this hypothesis, we systematically examined the functional role of ETC complex I in T-cell lymphoma. Our findings provide direct evidence that ETC complex I is integral to IL-17A-driven metabolic reprogramming, emphasizing its role in maintaining mitochondrial function and supporting tumor cell survival.

In line with these observations, analysis of blood and skin biopsies from CTCL patients revealed upregulated levels of ETC complex I. Similarly, elevated ETC complex I protein levels were observed in IL-17A-treated lymphoma cell lines. Functional studies further demonstrated that IL-17A-treated lymphoma cells exhibited enhanced ETC complex I activity, along with an increased NAD⁺/NADH and ATP/ADP ratio. By sustaining the cellular NAD^+^ pool and maintaining the NAD^+^/NADH ratio, ETC complex I supports the activity of mitochondrial malate dehydrogenase (MDH2), promoting aspartate production and cell proliferation (34). The high levels of ATP generated fulfill the elevated bioenergetic demands of malignant cells. Metabolic shifts favoring OXPHOS, leading to heightened ATP generation, have been observed in various advanced leukemias, lymphomas, and solid tumors (24,35,36).

Interestingly, IL-17A-treated lymphoma cells also showed a significant decrease in reactive oxygen species (ROS) and elevated mitochondrial membrane potential (MMP). The unexpected reduction in ROS may be attributed to ETC complex I-induced expression of ERK5 (extracellular signal-regulated kinase 5), which plays a crucial role in initiating antioxidant responses (37). While ROS levels are typically elevated in tumors to drive oncogenesis, excessive ROS can become toxic and trigger oxidative stress (38). Our data suggest that IL-17A helps maintain ROS at levels conducive to malignant T-cell survival in CTCL by mitigating oxidative stress through ETC complex I-mediated antioxidant responses. Moreover, the elevated MMP observed in IL-17A-treated cells aligns with reports linking high MMP in cancer cells to reduced apoptotic sensitivity (39). Collectively, our mechanistic studies reveal the tumor-promoting role of IL-17A through upregulation of ETC complex I, as evidenced by enhanced proliferation of malignant cells. These findings identify ETC complex I as a critical mediator of IL-17A signaling and underline its importance in CTCL pathogenesis. Supporting this, Kuntz et al. have also highlighted the essential role of OXPHOS in leukemia, further reinforcing the significance of ETC complex I in cancer progression (40).

Subsequent pharmacological inhibition of ETC complex I using mubritinib impaired its activity, leading to reduced ATP production, disrupted NAD⁺/NADH homeostasis, and increased ROS accumulation in lymphoma cells. Imbalanced NAD^+^/NADH levels cause ROS accumulation and dysregulate mitochondrial metabolism (41,42). These disruptions significantly curtailed cell proliferation, likely due to impaired mitochondrial metabolism and limited aspartate synthesis, critical for nucleotide biosynthesis during cell division (43). These findings further confirm the essential role of ETC complex I in supporting cancerous cell survival and proliferation.

Several studies have highlighted the potential of targeting OXPHOS as a therapeutic strategy in various disease settings (24,36,44,45). Specifically, OXPHOS has been shown as a metabolic vulnerability in therapy-resistant cancer cells, making it a promising target for novel treatments (21,40). Recently, a study on CTCL revealed that both OXPHOS and MYC are among the most enriched pathways in the disease and showed that disrupting these metabolic processes can inhibit lymphoma cell growth (46). Our study further elucidates the molecular mechanism and confirms that IL-17A plays a crucial role in modulating the OXPHOS in CTCL, and targeting ETC complex I may be a potential therapeutic strategy. In summary, IL-17A signaling induces mitochondrial metabolic rewiring via upregulating ETC complex I activity. Pharmacological perturbation of ETC complex I limits cell proliferation in lymphoma cell lines.

This study provides novel mechanistic insights into the IL-17A-dependent metabolic reprogramming of cancerous T-cells and identifies ETC complex I as a promising downstream therapeutic target for restricting malignant growth in T-cell lymphoma. By revealing the pathways involved in cellular homeostasis and their regulation, our findings contribute to a deeper understanding of metabolomics and its implications for medicine. Our study thus invites continued evaluation of targeting ETC complex I activity as a potential anti-cancer therapy, either alone or possibly in combination with existing therapies for CTCL. Furthermore, it paves the way for the development of additional targeted therapies that exploit altered mitochondrial metabolism in cancer cells.

## Author Contributions

Conceptualization: DA, BD, PS and RP

Methodology: DA and BD

Investigation: DA, BD, DM, VVS, SM, AB, DS and RLK

Visualization: DA, PS

Supervision: RP, PS

Clinical samples inclusion: HJ, TS, and NS

Writing—original draft: DA

Writing—review & editing: DA, BD, RP, HJ, PS

## Acknowledgments

We thank the individuals (CTCL patients and healthy volunteers) who donated the blood and skin biopsies for this study. We extend our gratitude to the family members of the patients, the medical staff of Tata Memorial Hospital, and the Central Super Resolution Confocal Microscopy, Laser Scanning Confocal Microscopy, and Spinning Disc Confocal Microscopy facility of IRCC, IIT Bombay, for their support. We acknowledge our lab member, Aalia Khan, at the IIT Bombay, for her assistance with the experiments. DA is thankful to the Council of Scientific & Industrial Research (CSIR) fellowship (09/087(1062)/2020-EMR-I) and the France Excellence Eiffel Fellowship (170882V). This work was supported by grants to R.P. from the Indian Council of Medical Research (ICMR) (RD/0119-ICMR000-001), Department of Biotechnology (DBT), Government of India, BT/INF/22/SP23026/2017 (Wadhwani Research Centre for Bioengineering, IIT Bombay).

We thank Dr. Michel Simon, Toulouse Institute for Infection and Inflammatory Diseases, for their constructive criticism and advice on this manuscript. This work was partially supported by grants to P.S. from the French National Research Agency (ANR) Young Investigator grant (ANR-21-CE12-0010), Cancer Research Foundation (ARCPJA22021050003683), La Ligue Foundation for Cancer (Haute-Garonne, 285458), Centre national de la recherche scientifique (CNRS), Université de Toulouse, Toulouse, France.

## Conflict of interest

The authors declare no potential conflict of interest

## Abbreviations

CTCL: Cutaneous T-cell Lymphoma
IL: Interleukin
ETC: Electron Transport Chain
MF: Mycosis Fungoides
SS: Sezary Syndrome
MMP: Mitochondrial Membrane Potential
ROS: Reactive Oxygen Species

## Notes

doi:10.25345/C5M902F58

### Competing Interest Statement

The authors have declared no competing interest.

