## Supplementary material for "IL-17A potentiates malignant T-cell viability via electron transport chain complex I in cutaneous T-cell lymphoma": Sup files

Diksha Attrish et al.

**This PDF file includes:**

Supplementary Text
Legends for Tables S1 to S2
Figures S1 to S5

**Other Supplementary Material for this manuscript includes the following:**

Table S1 to S2

**Supplementary Text**

**Supplementary Methods**

**Computational modeling of the MitoCore model**

We adopted a model-driven approach, utilizing Flux Balance Analysis (FBA) to quantitatively assess metabolic fluxes across cellular metabolic networks (1–3). Metabolic reactions were first encoded as a stoichiometric matrix (S) with dimensions [$m \times n$], where $m$ represents the number of metabolites and $n$ denotes the number of biochemical or transport reactions. In this matrix, negative coefficients indicate metabolite consumption, positive coefficients signify production, and a coefficient of zero denotes no involvement in the reaction. The model operates under the steady-state assumption, where the net production and consumption of each metabolite are balanced. Based on this assumption, constraints were calculated, and upper limits and lower limits were assigned to individual fluxes. To determine the most probable flux distribution, maximum ATP production was set as the objective function. These parameters were imported into the COBRApy environment and optimized using the built-in Gurobi linear program solver, and metabolic simulation was performed.

**Supplementary Figures and legends**

**Figure S1. (Related to Figure 1) CTCL patients exhibit higher expression of IL-17RA in CD3+ lymphocytes from skin biopsies.**

(**A**) Violin plot indicating the expression of IL-17RA in cutaneous T cell lymphoma (CTCL) and healthy control (HC) samples. **(B)** Dot plot showing the average IL-17RA expression level and percentage of T cells expressing IL-17RA of CTCL patients compared to healthy controls (HC). Data used for these analyses were from GSE128531.

**Figure S2. (Related to Figure 2) IL-17RA expression and dysregulated metabolic pathways due to IL-17A signaling.**

Representative histogram and quantification of CD25 (T-cell activation marker) **(A)** and IL-17RA **(B)** expression in healthy T-cells activated with CD3/CD28 dynabeads. (**C-D**) Representative histogram and quantification of IL-17RA expression in T-cell lymphoma cell lines **(C)** HuT 78 and **(D)** MJ. **(E)** Overview of multi-omics integrated analysis; Green arrows indicate the metabolic reactions with high flux as depicted by metabolism simulation analysis. APC: Allophycocyanin, PE: phycoerythrin. Values are expressed as mean ± SEM.

**Figure S3. (Related to Figure 3) CTCL patient samples exhibit higher expression of NDUFS2, and IL-17A upregulates ETC complex I expression.**

**(A)** Violin plots indicate the expression of NDUFS2 in CD3+ lymphocytes from skin biopsies of all cutaneous T-cell lymphoma (CTCL) and healthy control (HC) samples**. (B)** Dot plots showing the average expression of NDUFS2 in CTCL patients in comparison to healthy controls (HC). Data used for these analyses were from GSE128531. **(C-D)**. Histogram plot and statistical analysis quantifying ETC complex I expression in **(C)** MJ cells, **(D)** activated T-cells. Flow cytometry quantification of MTCO1 expression in **(E)** Jurkat cells, **(F)** HuT 78 cells, **(G)** MJ cells, and **(H)** activated T-cells. Values are expressed as mean ± SEM. UT: untreated, FITC: Fluorescein isothiocyanate, APC: Allophycocyanin.

**Figure S4. (Related to Figure 4) IL-17A affects mitochondrial metabolism and prevents apoptosis.**

Representative histogram and bar plot quantifying MMP in **(A)** Jurkat and **(B)** HuT 78 cells. **(C)** Flow cytometry dot plots and statistical line plots quantifying change in apoptotic cells due to IL-17A treatment in Jurkat and HuT 78 cells. Values are expressed as mean ± SEM. PE: phycoerythrin.

**Figure S5. (Related to Figure 5) Inhibiting ETC complex I perturbed MMP.**

Effect of increasing concentrations of mubritinib (50 nM to 1000 nM) on cell viability of **(A)** Jurkat and

**(B)** HuT 78 cells (n=3). Data are expressed as a percentage relative to UT (control). DMSO was used as vehicle control. **(C)** Flow cytometry dot plots and bar plot; Quantifying MMP upon mubritinib treatment (n=4) in HuT 78 cells. Values are expressed as mean ± SEM. $^*p < 0.05$, $^{**}p < 0.01$, $^{***}p < 0.001$ (Student paired t test). US: unstained, UT: untreated, DMSO: dimethyl sulfoxide (vehicle control), FITC: Fluorescein isothiocyanate.

**Tables S1-S2 in a separate file**

**Table S1. List of significantly differentially expressed proteins in Jurkat cells in response to IL-17A treatment.** Significantly differentially expressed proteins obtained from the proteome discoverer after applying the following filters: Protein False detection rate (FDR) confidence: 0.01, abundance ratio ≥ 1.2 and ≤ 0.667, and p value ≤ 0.1.

**Table S2. List of significantly differentially detected metabolites and their concentrations.** Catalog of significantly identified metabolites and their concentrations derived from NMR spectra analysis using Chenomx NMR suite 8.3.

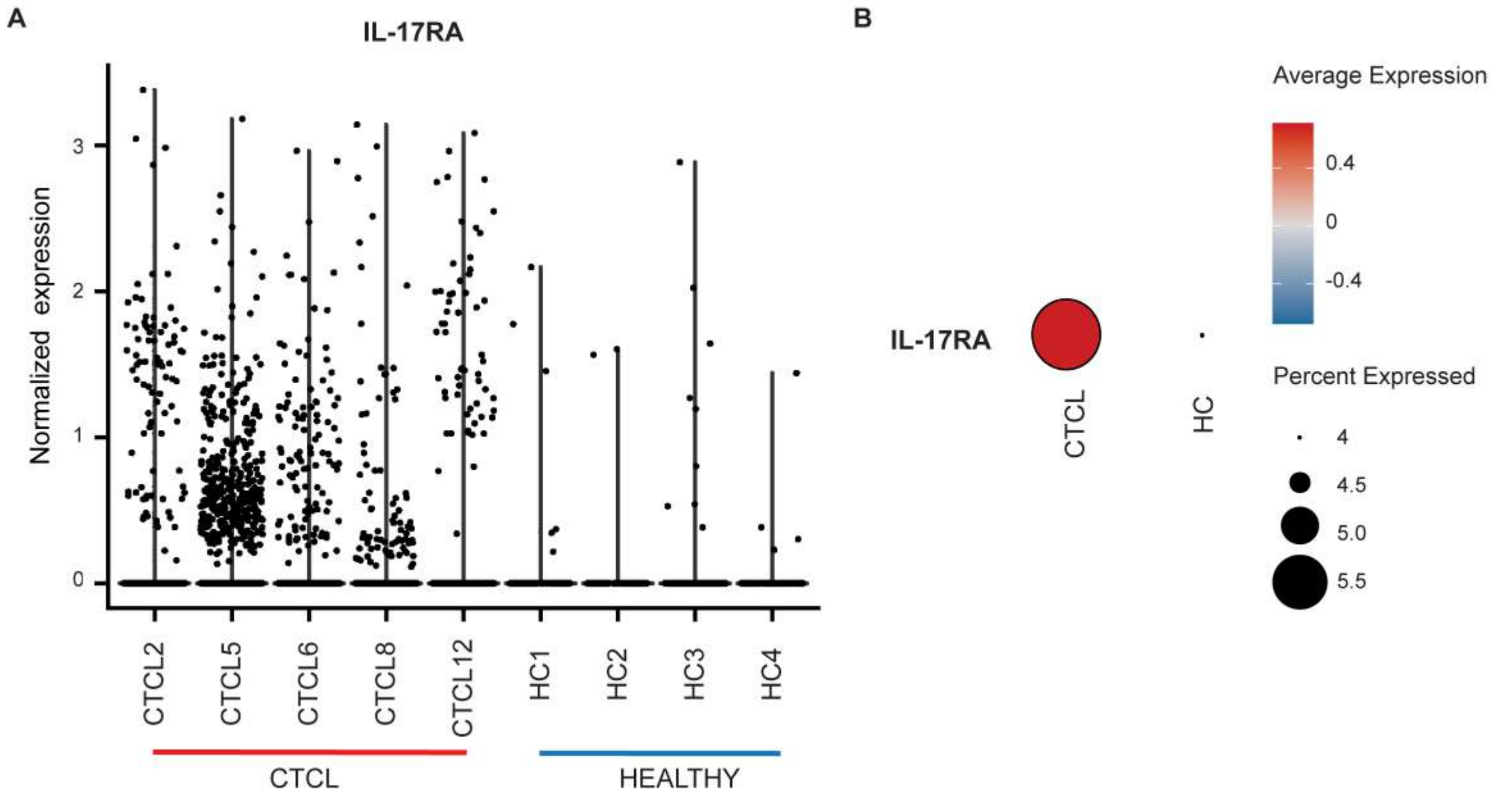


**Attrish et al., Figure S1**

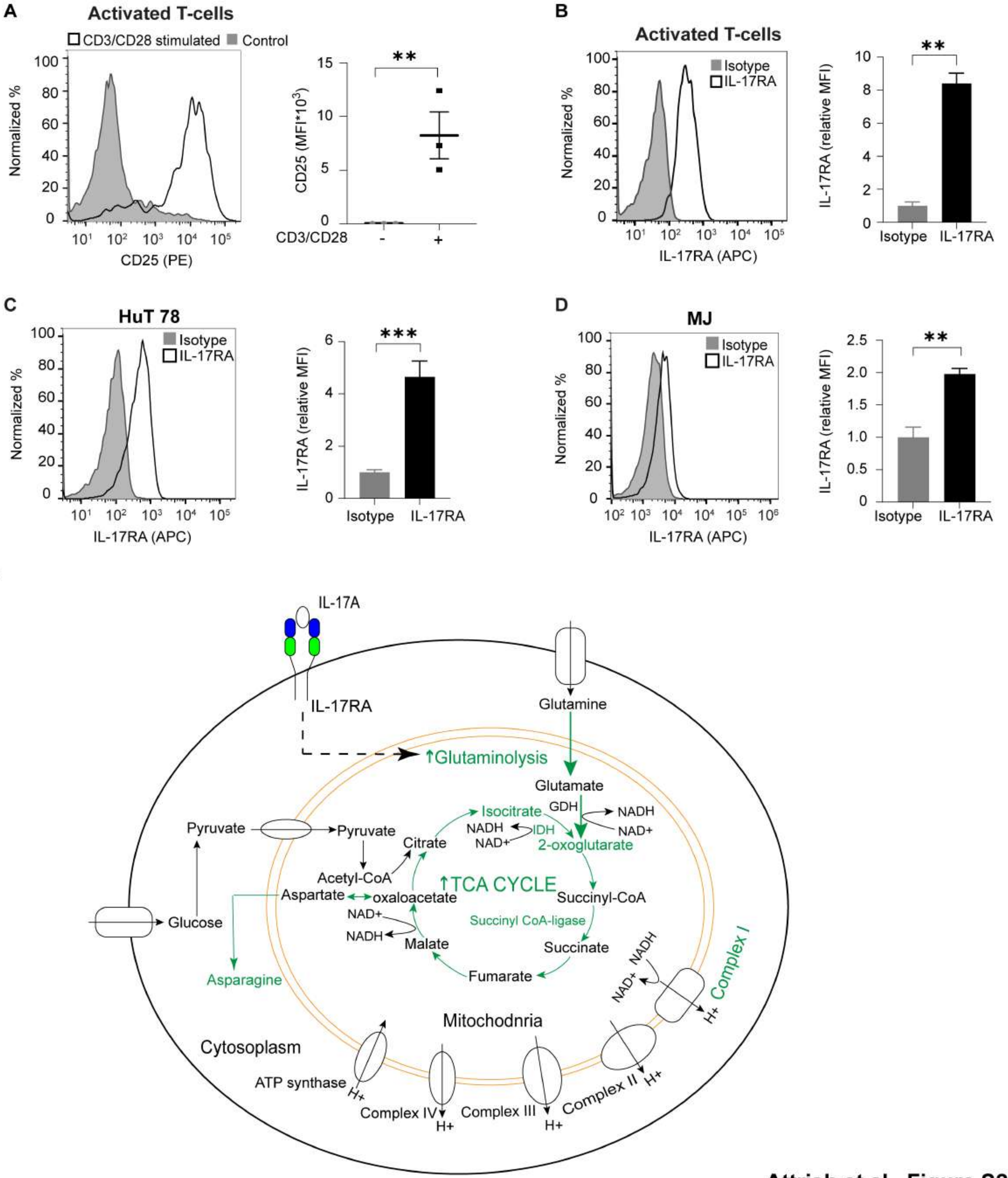


Attrish et al., Figure S2

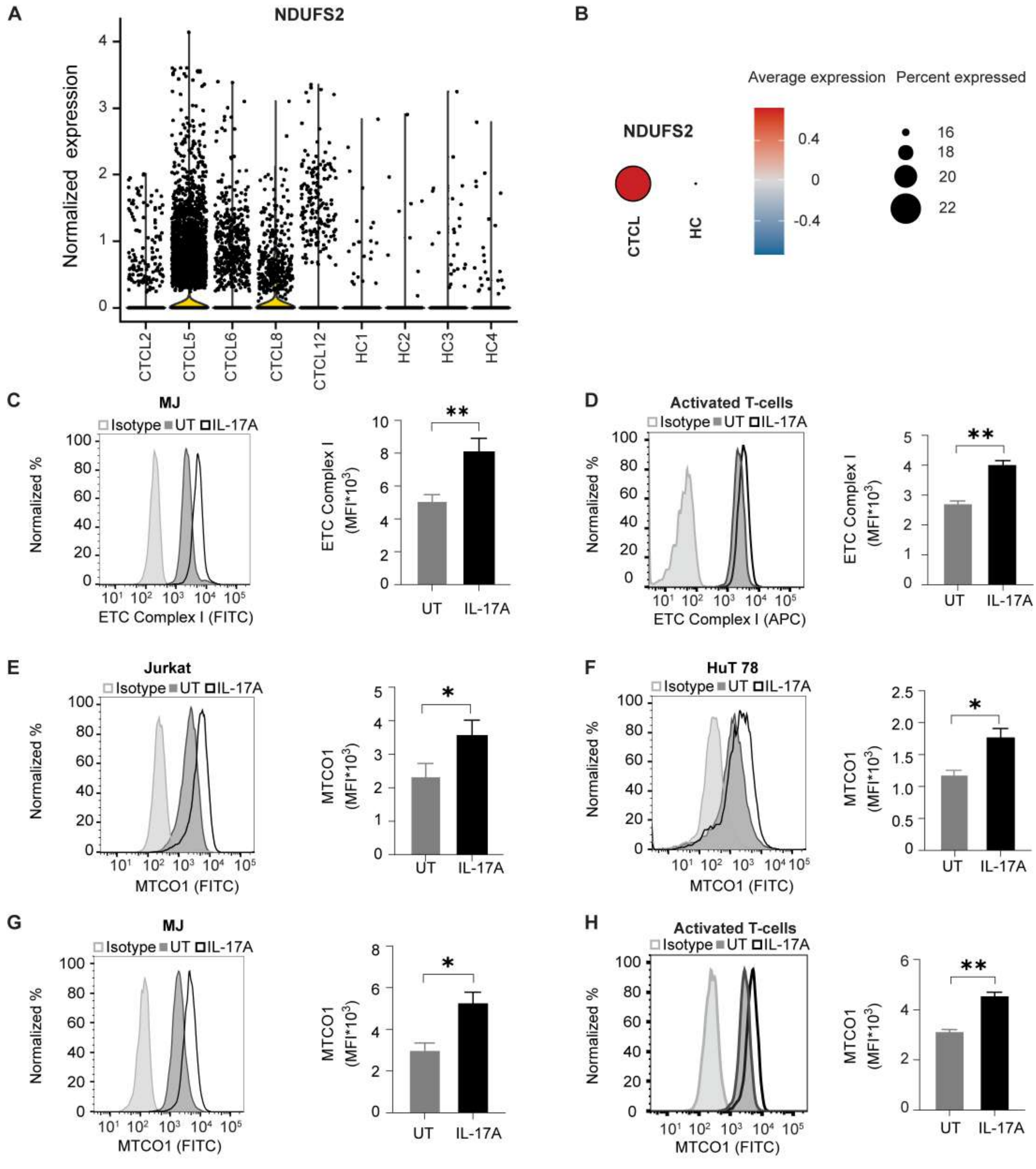


Attrish et al., Figure S3

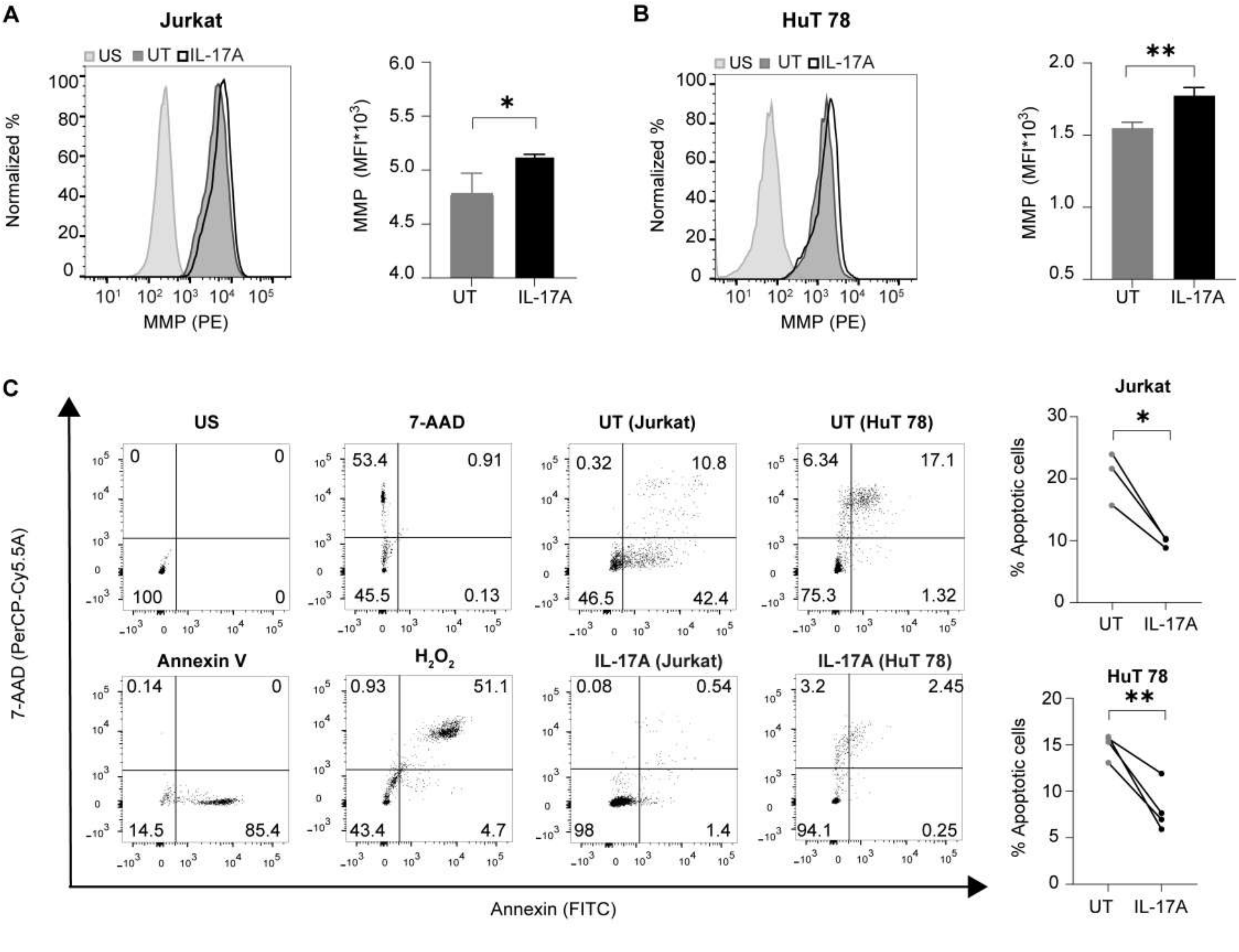


**Attrish et al., Figure S4**

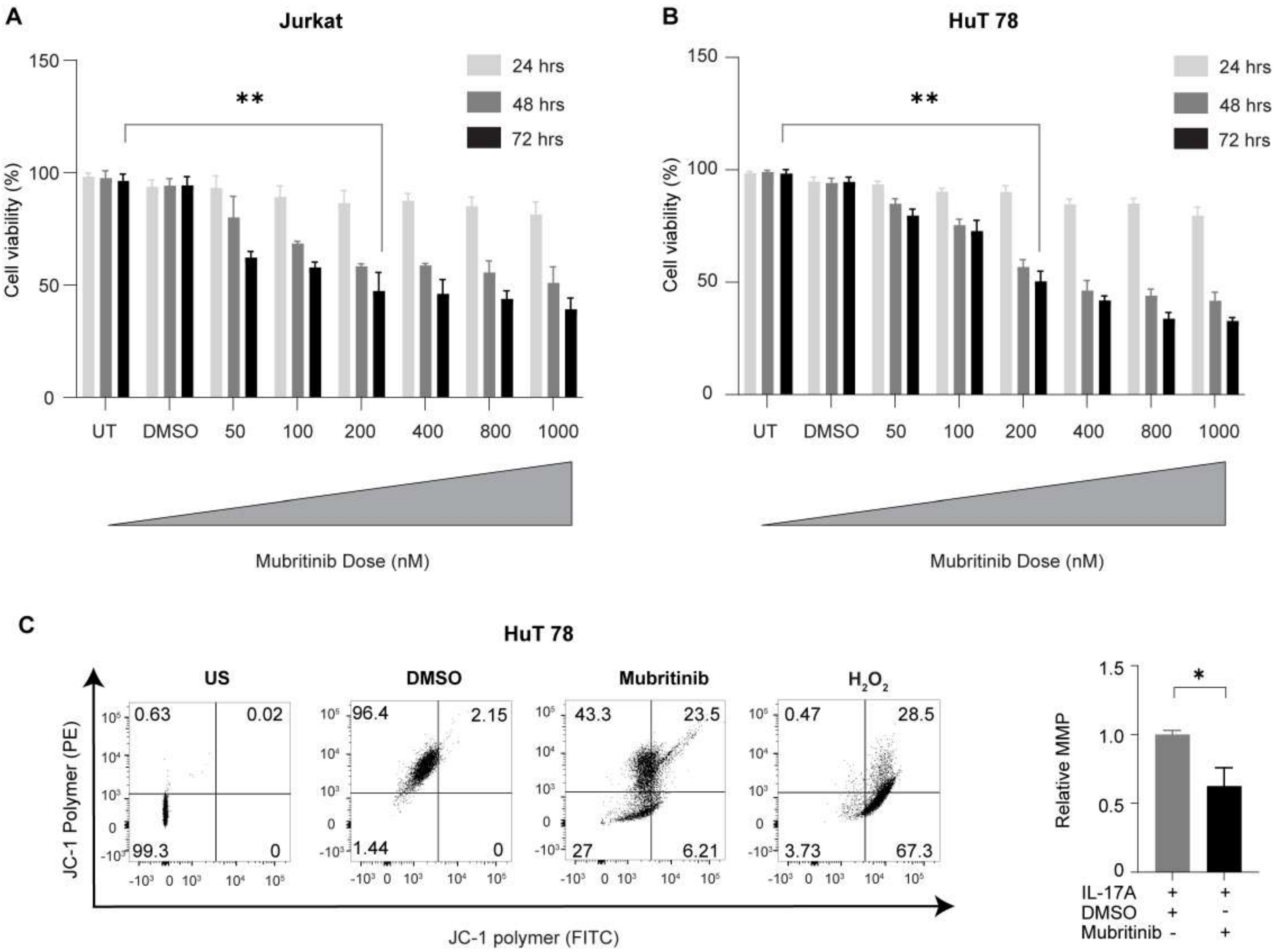


**Attrish et al., Figure S5**
